# Dibutyryl cyclic AMP downregulates tenascin-C in neurons and astrocytes and reduces AAV-mediated gene expression in DRG neurons

**DOI:** 10.1101/2025.05.13.653846

**Authors:** Katerina Stepankova, Anda Cimpean, Barbora Smejkalova, Radovan Holota, Lenka Bachanova, Jiri Cerny, Vojtech Sprincl, Dana Marekova, Joëlle van den Herik, Fred de Winter, Pavla Jendelova, Lucia Machova Urdzikova

**Author notes:** KS and LMU share corresponding authorship Katerina Stepankova, Lucia Machova Urdzikova.

## Abstract

Functional recovery after spinal cord injury (SCI) is hindered by the limited ability of axons to regenerate in the adult mammalian central nervous system (CNS). Overcoming this barrier is critical for achieving effective recovery. Axonal regeneration depends on the activation of intracellular processes like transcription factor induction, protein and lipid trafficking, and cytoskeletal remodelling. Targeting these pathways offers a promising approach for promoting neuronal repair. This study examined the combined therapeutic effects of dibutyryl-cAMP (db-cAMP), which primes neurons for growth, and integrin α9 overexpression, which supports axonal extension. Using in vitro models with dorsal root ganglion (DRG) neurons and astrocytes, as well as an in vivo SCI model, we evaluated the potential of this approach. In vitro, the combination of db-cAMP and integrin α9 significantly enhanced neuronal growth. However, in vivo results were less consistent, with db-cAMP affecting AAV-mediated transcription and the expression of tenascin C (TnC) in neurons and astrocytes. These findings highlight the potential of modulating intracellular signalling and integrin activation but underscore the challenges posed by the complexity of the in vivo environment. Further studies are necessary to unravel these mechanisms and refine therapeutic strategies for effective SCI recovery.

**Graphical abstract:** 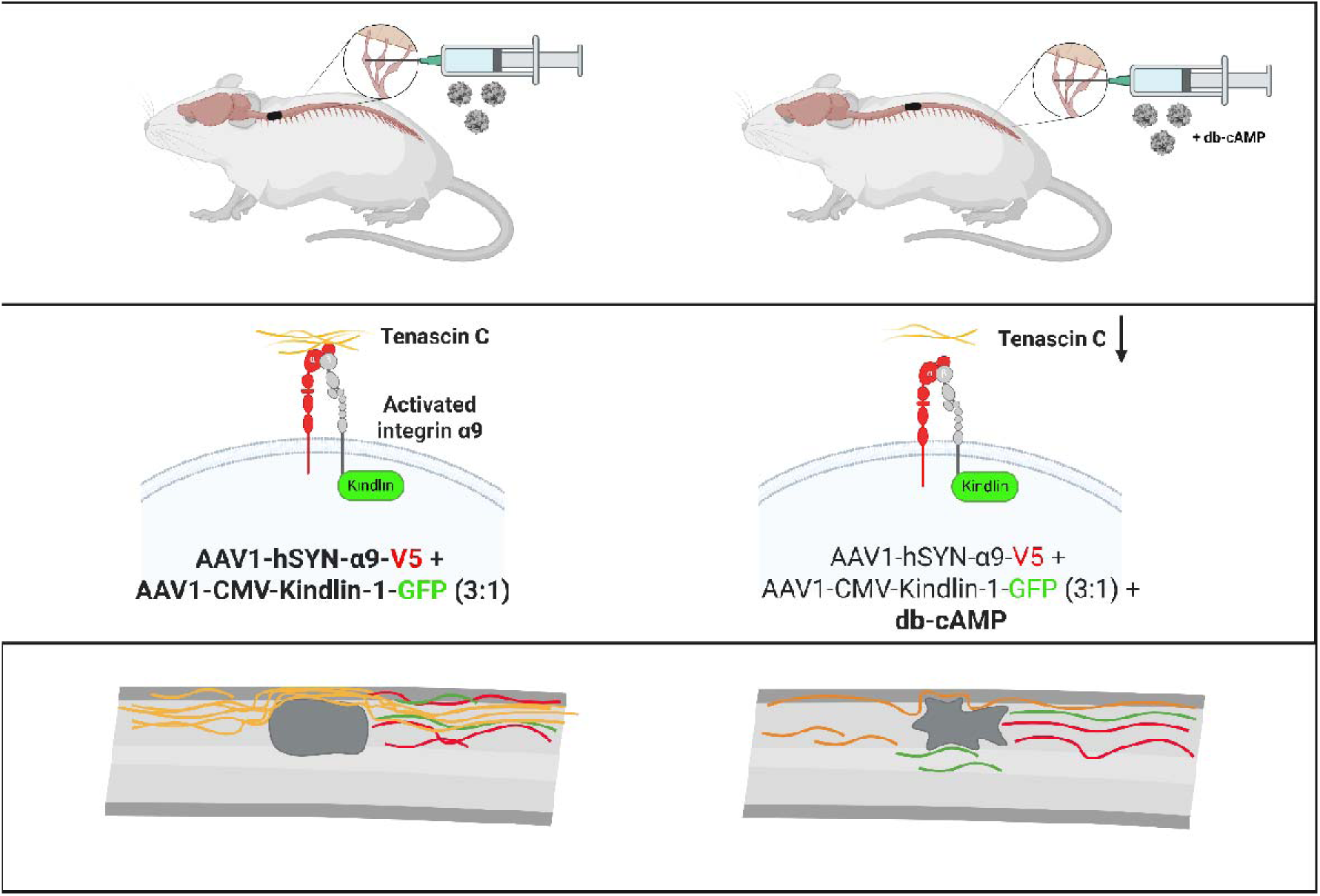

## Introduction

Functional recovery after spinal cord injury (SCI) depends on the ability of neurons to regenerate^1^. However, in the adult mammalian central nervous system (CNS), most injured axons fail to regenerate, representing a significant barrier to recovery after CNS trauma^1,2^. Successful axonal regeneration requires the activation of several intracellular processes, including the induction of transcription factors that initiate regeneration programmes, the transport of essential lipids and proteins, and the reorganisation of the cytoskeleton to form growth cones^3,4^. However, insights from developmental stages provide potential strategies to promote regeneration by exploiting the molecular pathways involved in axon growth^5^.

Axon regrowth after SCI is severely limited in the adult mammalian CNS, resulting in permanent sensory and motor impairments^1,6^. To uncover the molecular pathways and key regulators that facilitate axon regeneration, researchers have explored several approaches^2–4^. These include studying developmental mechanisms associated with axon growth, conducting molecular screens in invertebrates and non-mammalian vertebrates, and studying injury-induced responses such as the widely studied conditioning lesion paradigm^2–6^.

Axon regeneration after a conditioning lesion is effectively demonstrated in sensory dorsal root ganglia (DRG) neurons^7^. The pseudo-unipolar structure of DRG neurons allows them to transmit afferent information from the periphery to the spinal cord and supraspinal regions^8^. The regenerative potential of DRG neurons after SCI can be enhanced by prior activation of intrinsic growth mechanisms, such as conditioning^9^. This approach, identified in the late 20th century, has since become a key model for the study of sensory neuron regeneration and has contributed to the discovery of several regenerative signalling pathways^10^. Increasing intracellular levels of cyclic AMP (cAMP) has emerged as one of the most successful strategies for promoting axonal regeneration in the CNS^11^. Intracellular cAMP can be increased by various methods, including peripheral conditioning lesions, administration of cAMP analogues (such as db-cAMP), and priming with neurotrophins. Despite its ability to prime neurons for regeneration, the extent of axonal growth induced by db-cAMP conditioning remains limited, particularly in the spinal cord where even primed sensory neurons face challenges in extending axons beyond the injury site^12,13^.

Recent studies have shown that viral vector-mediated delivery of integrin α9 to DRG neurons, in combination with activation by kindlin-1, significantly enhances sensory axon regeneration in the CNS^14–16^. Integrins, a family of heterodimeric cell surface receptors, are essential for linking the extracellular matrix (ECM) to the cytoskeleton and intracellular signalling pathways to support processes such as adhesion, migration and neurite outgrowth^17^. Among these, integrin α9 plays a critical role in maintaining axon growth^18^. It interacts with tenascin C (TnC), the primary ligand for integrin α9 in the CNS^19^. TnC is an ECM protein that is expressed during development and in neurogenic regions of the adult CNS and is upregulated following CNS injury, particularly around the lesion border^20–24^. By binding to integrin α9, TnC promotes regeneration^14^. However, mature CNS neurons typically lack integrin α9 expression, limiting their regenerative potential^14^. When activated by kindlin-1^25^, integrin α9 enables DRG neurons to overcome inhibitory environments and promotes increased axonal growth in the adult CNS after SCI when its expression is upregulated using viral vectors^16^.

The aim of this study was to determine whether the combination of db-cAMP priming and integrin α9 overexpression could enhance axon regeneration after spinal cord injury (SCI). Db-cAMP primed neurons for immediate growth, while integrin α9 provided structural support to overcome the inhibitory barriers of the CNS. In vitro experiments with DRG neurons tested the hypothesis that activated integrin α9 promotes neurite outgrowth under inhibitory conditions. The results showed that AAV-mediated overexpression of integrin α9 in combination with db-cAMP significantly improved neurite outgrowth. Based on these findings, we moved to an in vivo SCI model to assess whether db-cAMP could enhance the regenerative potential of integrin α9 overexpression in DRG neurons following SCI in rats.

Mechanistically, we found that while db-cAMP and AAV-mediated integrin overexpression can modulate the growth state of DRG neurons in culture, the synergistic effects observed in vitro do not fully translate to the in vivo environment. In addition, db-cAMP appears to affect both AAV-mediated transcriptional efficiency and intrinsic expression of TnC, affecting not only DRG neurons but also spinal astrocytes. Remarkably, changes in TnC expression following db-cAMP application have been observed in rodents and appear to represent a conserved mechanism within the mammalian CNS.

## Results

### Under both permissive and inhibitory conditions in vitro, db-cAMP treatment increases neurite length and neurite biogenesis

TnC and Aggrecan (Agg) are two major extracellular proteins that are inhibitors of neurite outgrowth^26,27^. However, axon outgrowth is possible when axons express activated integrin α9 (α9 integrin activated via kindlin-1, α9-K1), which uses TnC as substrate^14,15^. We first validated the previous findings on the mechanisms and effects of the α9 integrin and its activator, kindlin-1, on neurite length in adult DRG neurons under permissive (coverslips coated with PDL only) and inhibitory conditions (coverslips coated with TnC and Agg) (Figure 1 A). Under permissive conditions, neurons transduced with α9 or α9-K1 exhibited significantly longer neurites (450 µm and 500 µm, respectively) compared to non-transduced (NT) or GFP-transduced neurons (180 µm and 160 µm, respectively). This was indicative of enhanced neurite outgrowth. No significant differences were observed between the neurite lengths of the α9 transduced neurons and the α9-K1 transduced neurons (Figure 1 B). When the cells were cultured in an inhibitory environment, both NT and GFP-transduced cells had significantly reduced neurite lengths. The average length was approximately 80 µm. Both α9 and α9-K1 transduced cells were able to overcome this inhibition and grew longer neurites. However, α9-K1 transduced cells showed a significantly higher average neurite length of 350 µm compared to 210 µm in the α9 integrin transduced cells (Figure 1 C). These results confirm previous findings that α9 integrin can use inhibitory TnC to promote regeneration. Furthermore, the intracellular activator Kindlin-1 further enhances integrin-driven neurite outgrowth, even in the presence of Agg.

**Figure 1.**
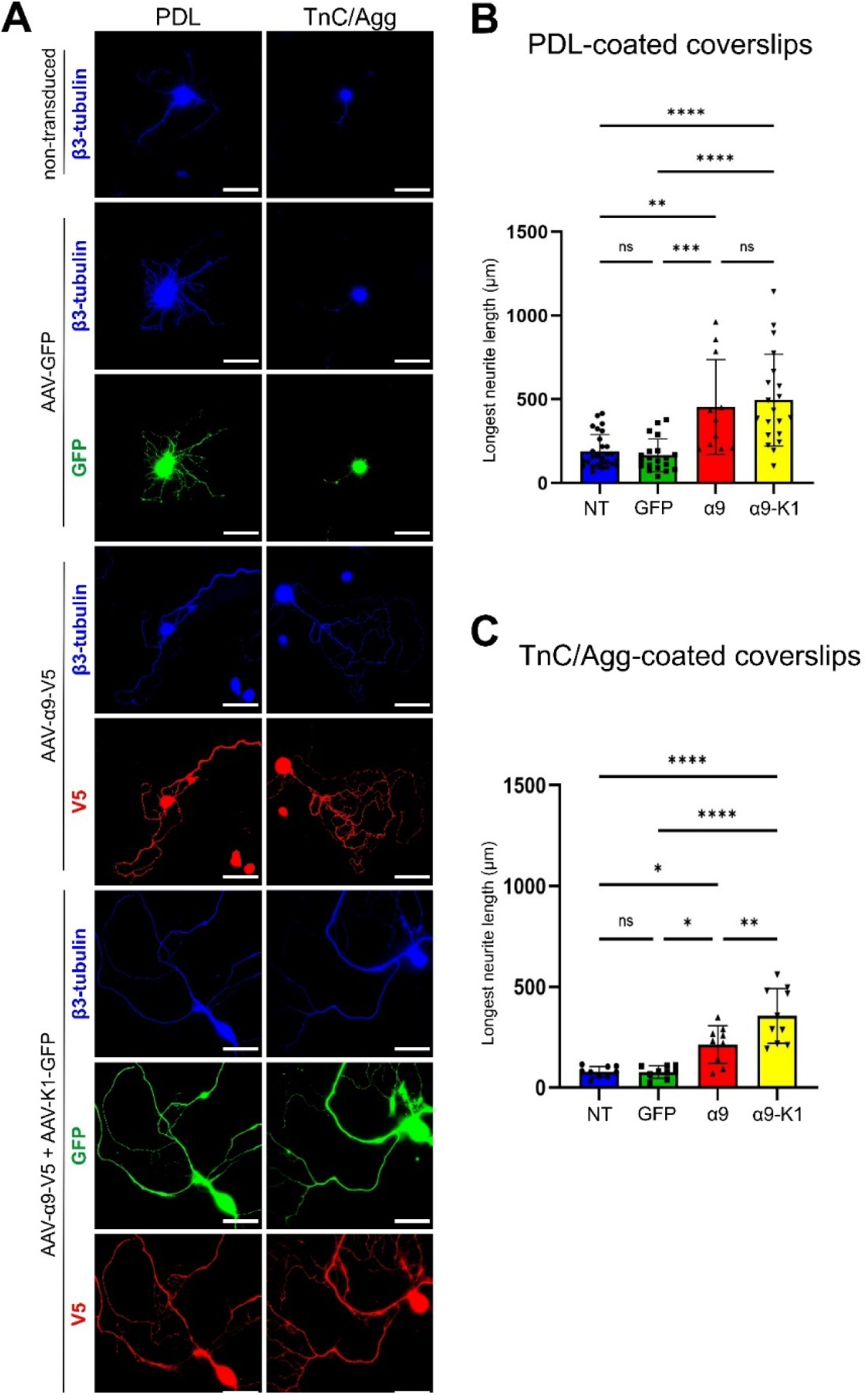
Activated integrin **α**9 promoted neurite outgrowth under permissive and inhibitory conditions. (**A**) Representative images of Non-, GFP-, α9 integrin- and α9 integrin kindlin-1 transduced neurons growing on PDL- and Agg/TnC-coated coverslips. Quantification of (A). The longest neurite (µm) growing on PDL-coated coverslips (**B**) and on Agg/TnC-coated coverslips (**C**). The longest neurite of each neuron was quantified from at least three separate experiments. The individual values represent single neurites. Statistical analysis was performed by one-way ANOVA. ns >0.5, *p < 0.05, ** p < 0.01, *** p < 0.001, **** p < 0.0001. Data are presented as mean ± SEM. Scale bar: 70 µm.

Having confirmed that our experimental design was consistent with the literature, we examined the effect of db-cAMP treatment on neurite length. NT, GFP-, α9- and α9-K1-were treated with 2 mM db-cAMP throughout the culture period (5 DIV), then fixed and neurite lengths compared with untreated conditions (Figure 2 A). On PDL-coated coverslips, db-cAMP treatment significantly increased neurite length in NT and GFP-transduced neurons from 170µm to 250µm on average (Figure 2 B, C). Neurite length in α9 and α9-K1 transduced cells was not significantly affected by db-cAMP treatment (p=0.1 and p=0.8, respectively) (Figure 2 D, E). Neurite outgrowth was then tested in the presence of TnC and the inhibitory CSPG Agg. On TnC and Agg-coated coverslips, cAMP treatment resulted in longer neurites in NT and GFP-transduced neurons. Neurite length increased from 78 µm to 160 µm (Figure 2F, G). Of note, we also observed a significant increase in neurite outgrowth in α9-transfected neurons (from 200µm to 325µm) and α9-K1-transfected neurons (from 320µm to 510µm) in the presence of cAMP (Figure 2 H, I). Interestingly, while integrin overexpression led to a less complex neuronal morphology—reflected by a reduced total number of neurites per neuron—cAMP treatment conferred an additional, unexpected benefit. On both PDL- and TnC/Agg-coated coverslips, the number of cells bearing neurites doubled under cAMP-treated conditions compared to untreated controls, suggesting that cAMP may help counterbalance the morphological simplification associated with integrin overexpression (Figure 2 L, M). (Figure 2 L, M). The total number of cells was not significantly different between treated and untreated (Figure 2J, K), indicating that the effect of cAMP was on increasing neurite biogenesis rather than cell adhesion or survival. These results suggest an additional beneficial role for cAMP in promoting neurite biogenesis, corroborating previous reports of the positive effect of cAMP on neurite length under both permissive and inhibitory conditions.

**Figure 2.**
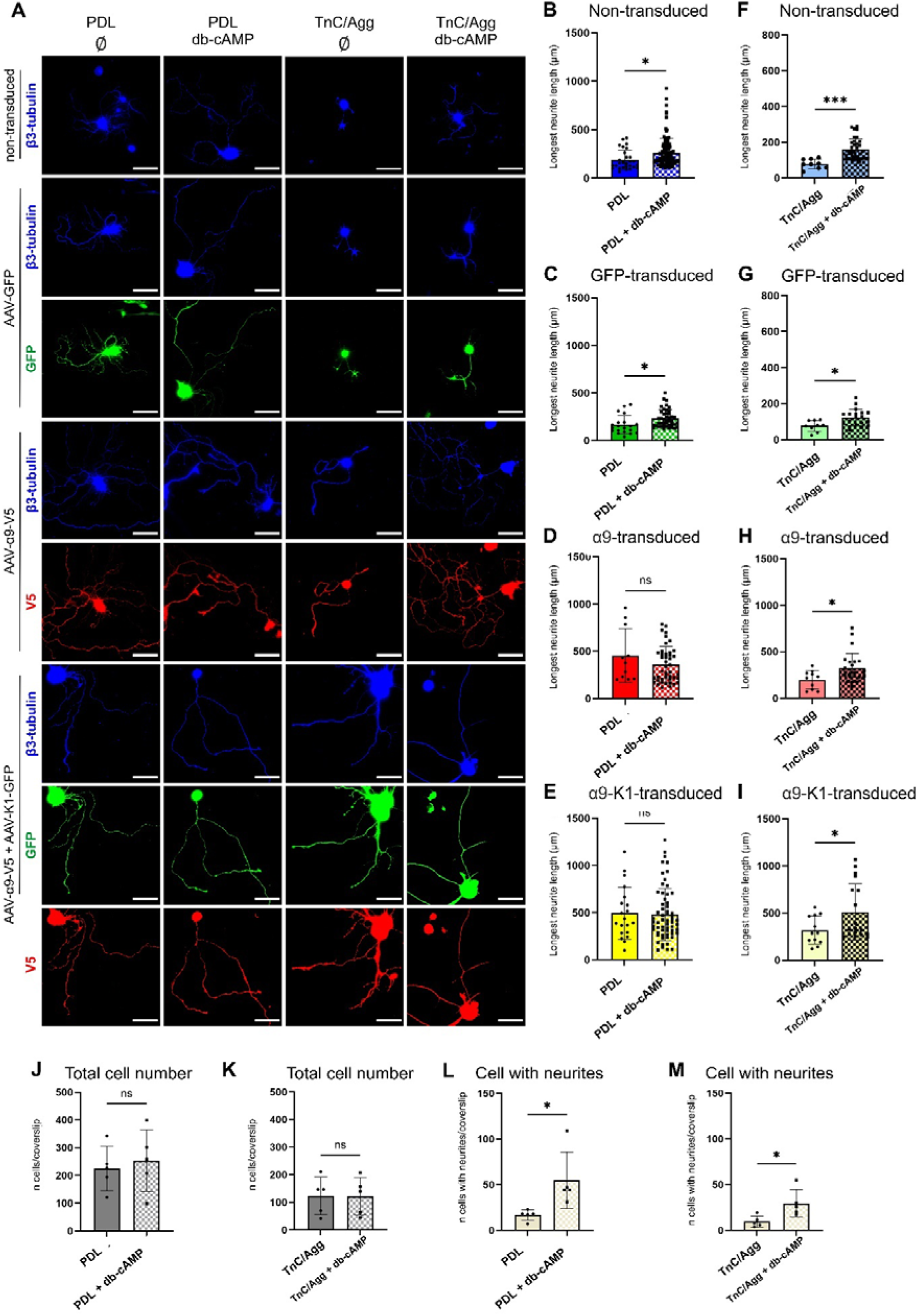
Db-cAMP treatment enhanced neurite length and promoted neurite formation under both permissive and inhibitory *in vitro* conditions. (A) Representative images of Non-, GFP-, α9-integrin- and α9-integrin Kindlin-1-transduced neurons growing on PDL- and Agg/TnC-coated coverslips treated with or without db-cAMP. Under permissive *in vitro* conditions (PDL-coated coverslips), db-cAMP significantly increases neurite length in non-(B) and GFP-transduced (C) cells but does not affect neurite length in α9-(D) and α9-K1-transduced (E) cells. Under inhibitory conditions (Agg/TnC-coated coverslips), the same effect of db-cAMP on neurite outgrowth was observed in non-(F) and GFP-transduced (G) cells. In contrast to permissive conditions, under inhibitory conditions *in vitro*, db-cAMP also increases neurite length in α9 (H) and α9-K1 (I) transduced cells. (J, K) Bar graphs show that there was no significant difference between the total number of cells in the absence or presence of db-cAMP. However, a significant increase in the number of neurite-bearing cells was observed in the presence of db-cAMP on both PDL- and Agg/TnC-coated coverslips. Quantification was performed from at least three separate experiments. In (B-I) the individual values represent single neurites and in (J-M) the individual values represent separate experiments. Statistical analysis was performed by unpaired t-test. ns >0.5, *p < 0.05, *** p < 0.001. Data are presented as mean ± SEM.

### Regeneration after SCI is not enhanced by the combination of activated integrin α9 and db-cAMP

In our *in vitro* setup, we assessed the immediate cellular responses and effects of db-cAMP treatment on neurite outgrowth and neuronal signalling. This approach allowed for precise control of experimental conditions and rapid evaluation of initial outcomes. To translate the positive effects observed *in vitro* to an *in vivo* context, we performed a dorsal column crush at the T10 level in rats, concurrently injecting the DRGs with either AAVs alone (AAV1-hSYN-GFP or AAV1-hSYN-α9-V5 + AAV1-CMV-K1-GFP) or with AAVs combined with db-cAMP at a final concentration of 10 mM.

To assess the potential therapeutic benefits of combining activated integrin and db-cAMP, we conducted a comprehensive battery of behavioural tests, including the BBB, Von Frey, plantar, and tape removal tests. Our results did not indicate that the combination of db-cAMP with a9-K1 was more effective in promoting axon regeneration following spinal cord injury (SCI) than a9-K1 alone. This study confirmed the findings of a previous study^16^, which demonstrated regeneration through the overexpression of activated α9 in DRG neurons after SCI (Figure 3 A, B, C, D, E).

**Figure 3.**
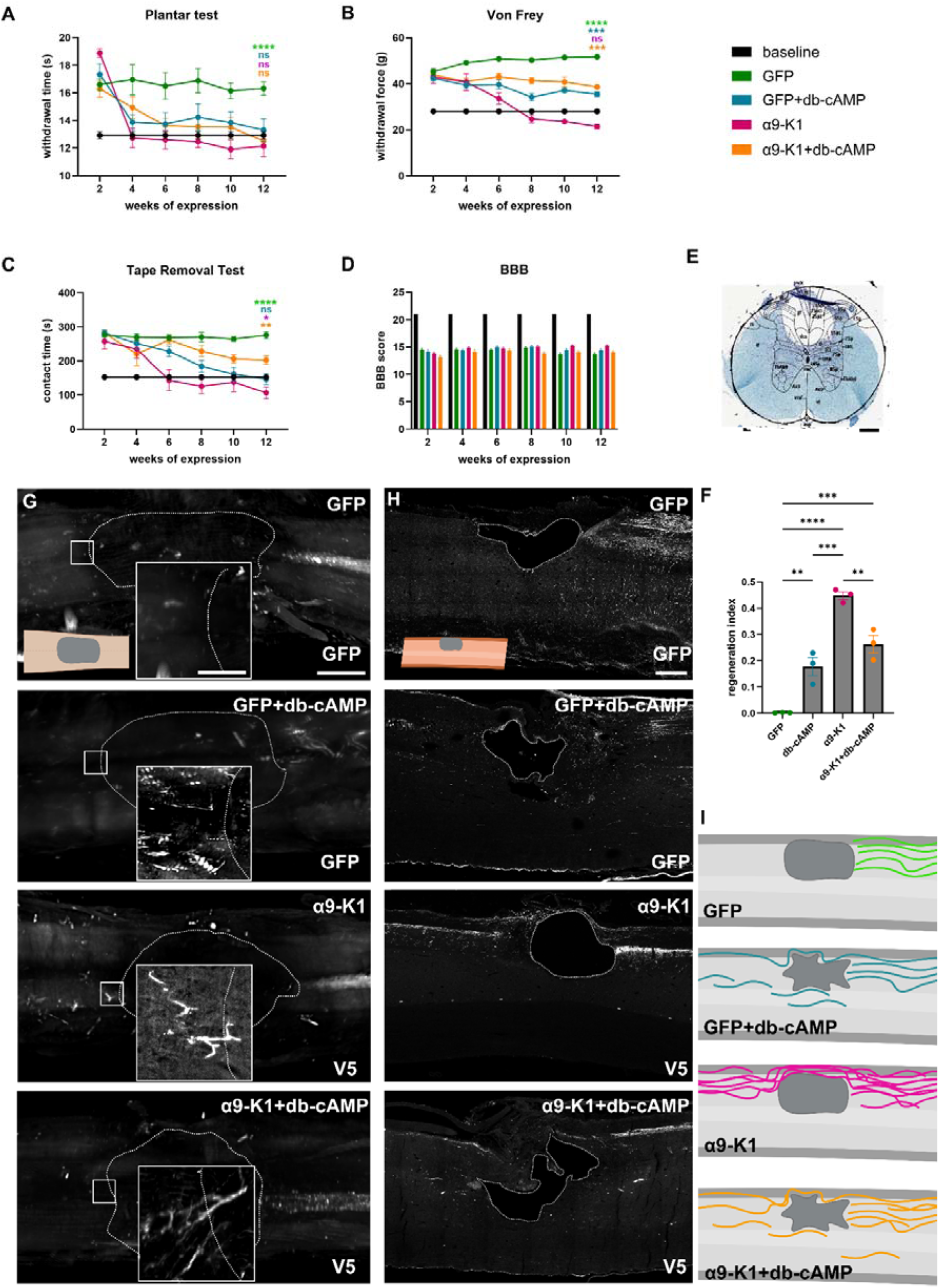
The combination of db-cAMP with activated integrin. α**9 does not provide additional benefits for axon regeneration or recovery compared to activated integrin** α**9 alone.** Bar graphs illustrate behavioural test results for evaluating three therapeutic approaches: (B) The plantar test (thermal sensation) shows all treated groups returning to baseline, outperforming the GFP control group. (**B**) The von Frey test (mechanical sensitivity) indicates sensory recovery only in animals overexpressing activated integrin. Animals treated with db-cAMP, with either AAV1-hSYN-GFP or AAV1-hSYN-α9-V5 + AAV1-CMV-K1-GFP, exhibited similar effects. (**C**) The tape removal test highlights the greatest improvement in the α9-K1 group, followed by GFP+db-cAMP. (**D**) The BBB test (locomotor abilities) reveals no significant differences, confirming consistent lesion characteristics. Behavioural data (mean ± SEM; n=6–12) are shown for the 12th week, with significance marked (*p_J<_J0.05, **p_J<_J0.01, ***p_J<_J0.001, ****p_J<_J0.0001; two-way ANOVA with Dunnett’s test). Full statistics and raw data are in the Supplementary Material. (**E**) Luxol Fast Blue-stained spinal sections, with drawings of ascending/descending tracts, show a regeneration index (ratio of axons above and below the lesion) in (F) (mean ± SEM; **p_J<_J0.01, ***p_J<_J0.001, ****p_J<_J0.0001; one-way ANOVA, Tukey’s test; n=3). (**G**) Light sheet images (scale bars: 200 µm, insets: 20 µm) highlight axons above the lesion, with lesion borders marked by dotted lines. (**H**) Confocal images of sagittal spinal sections (scale bar: 200 µm) show axon regrowth across groups, with orientation diagrams created using BioRender.com. (**I**) A schematic summary of findings highlights distinct differences between experimental groups. In GFP-treated animals, axons halted at the lesion edge. GFP+db-cAMP axons crossed the lesion, though not solely via the connective tissue bridge, with some below the lesion. In the α9+K1 group, axons grew upward along the connective tissue bridge. However, in the α9+K1+db-cAMP group, bridge usage was reduced. Additionally, db-cAMP altered lesion shape regardless of viral vector treatment.

In the current study, we observed minimal differences in outcomes between db-cAMP administered with either the GFP-containing vector or the a9-K1 vector mixture, suggesting that the choice of vector does not play a crucial role in regeneration in this context. In the plantar test, no significant differences were observed between the three treated groups and the baseline values, except that the GFP (control group) remained significantly less sensitive to thermal stimuli compared to the baseline after 12 weeks of testing. These findings suggest the effectiveness of an activated integrin and both db-cAMP groups in restoring the spinal warm-sensing pathways^28^ following dorsal column crush.

In the Von Frey test, only the a9-K1 group showed improvement, returning to baseline levels. This further confirms the potential of activated integrin in reconstructing sensory pathways. However, none of the animals in the db-cAMP or GFP groups reached baseline values, suggesting limitations in reconstructing long-projecting axons responsible for transmitting mechanical signals to the medulla^29^.

Lastly, in the tape removal test, we observed improvement in the GFP + db-cAMP group, which returned to baseline values. Additionally, animals in the a9-K1 group performed even better than baseline, indicating not only an ability to perceive the sticky tape on their paw but also some degree of learning. The a9-K1 + db-cAMP group showed improvement compared to the GFP control group but remained less sensitive to the tape than baseline values (Figure 3 A, B, C, D, E).

It is important to note that we did not include a sham control group in our animal study, as our primary focus was to evaluate the efficacy of three treatment groups against a single control group. This design minimized the number of animals used while still allowing for an effective assessment of the therapeutic effects of our interventions.

To further validate the behavioural test results, we used immunohistochemistry to visualize axons in sagittal sections of the spinal cord. The regeneration index was calculated as the ratio of axons counted above (+ 600 μm) versus below (−600 μm) the lesion site. Previously, we observed that axons with overexpressed activated integrin α9 achieved a regenerative index of approximately 0.5 at 600 µm above the lesion, extending as far as the medulla. A smaller number of axons reached 5 cm above the lesion, with a regenerative index of 0.19^16^. Axon distance travelled was not measured, as the combination therapy did not demonstrate a significant beneficial effect in this regard. The regeneration index results showed no axons above the lesion in the GFP control group (0.0022 ± 0.0007), while GFP+db-cAMP (0.18 ± 0.035), α9-K1 (0.45 ± 0.015), and the combination α9-K1+db-cAMP group (0.26 ± 0.034) showed different degrees of regeneration. An important aspect of the present results is that we successfully demonstrated the replicability of previously observed α9-mediated regeneration after dorsal column crush, yielding similar data. These findings are consistent with the behavioural outcomes, reinforcing the observed functional improvements. The regeneration index was assessed in spinal cords from three randomly selected animals per group, revealing clear differences between treatments (Figure 3 F, G).

Taken together, the results from the plantar test and the regeneration index suggest that thermal sensation improvement occurred across all three experimental groups, even though only a limited number of axons successfully crossed the lesion. This indicates that mechanisms beyond direct axonal regeneration, such as intraspinal circuit adaptations, likely play a crucial role in mediating thermal sensation recovery after SCI^30^.

### db-cAMP influences the efficiency of the AAV vector at a transcriptional level when administered concurrently with injections to the DRG

An interesting observation was that animals injected with db-cAMP showed comparable outcomes regardless of whether they received just AAV1-hSYN-GFP viral vector or in combination with treatment (AAV1-hSYN-a9-V5 + AAV1-CMV-K1-GFP; 3:1). This raised the question of whether db-cAMP influences viral expression when co-administered with these vector(s). To investigate this, we stained cleared DRG samples with GFP or V5 antibodies, visualised them using lightsheet microscopy, and then quantified the number of GFP- or V5-positive cells in Imaris software (Oxford Instruments, Abingdon, UK). We found no significant difference in the number of GFP or V5-positive cells between db-cAMP-treated and untreated groups (Figure 4 A, B, C).

**Figure 4.**
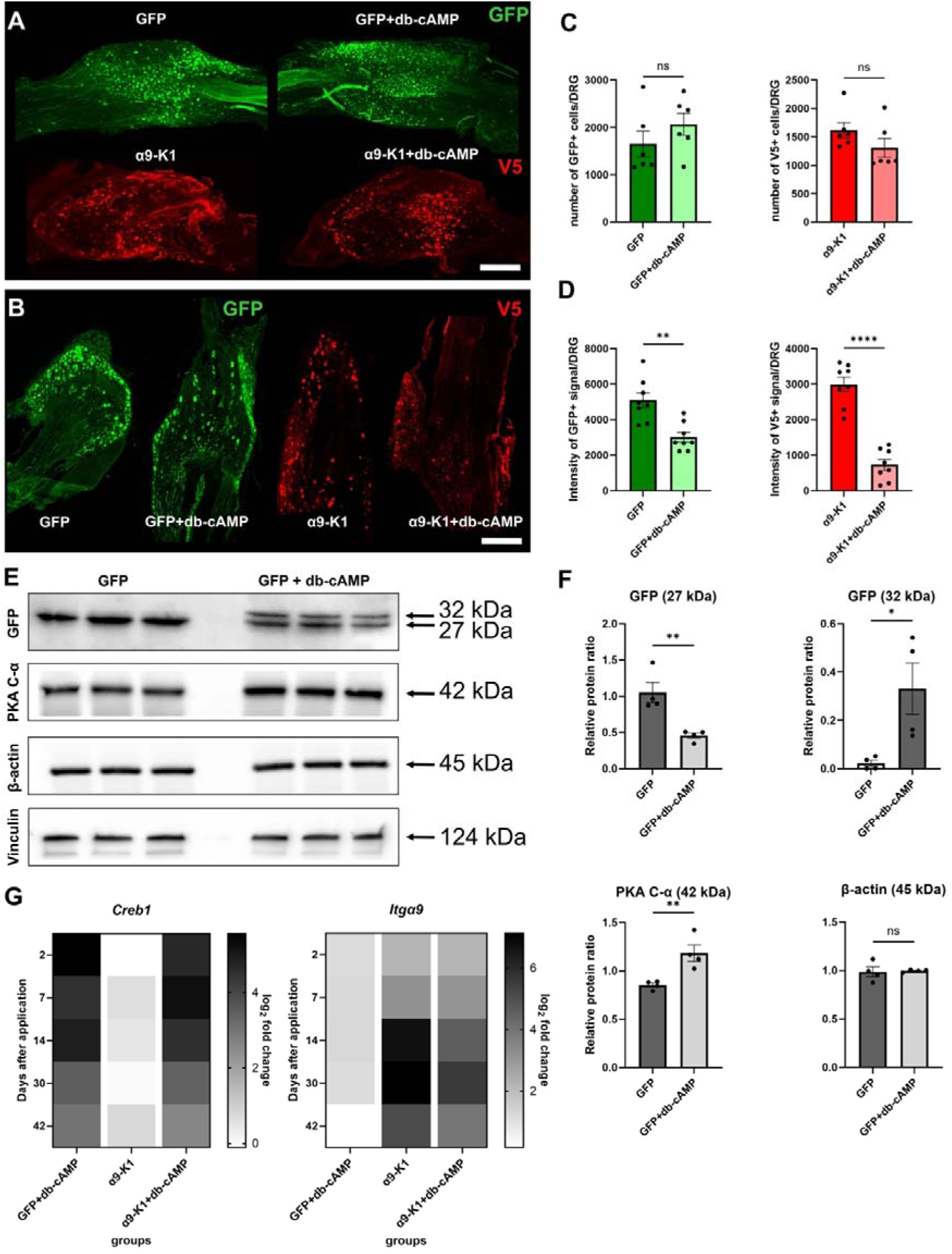
Db-cAMP influences the transcriptional efficiency of the AAV vector when co-injected into the DRG. (**A**) Representative snapshots from Imaris software of DRGs visualized using the Zeiss Lightsheet Z.1 following ECi clearing. Scale bar: 400 µm. (**B**) Representative confocal images of DRG sections showing varying signal intensities in the presence and absence of db-cAMP. For intensity measurements, all stained sections were imaged using the same laser power, gain, and airy units. Scale bar: 400 µm. (**C**) Quantification of (A). Bar graphs display the total number of GFP-positive or V5-positive cells in the DRG. Data are presented as mean ± SEM, analysed by unpaired t-test, ns > 0.5, *p < 0.05, n = 6 DRGs/group. (**D**) Quantification of (B). Bar graphs show the total GFP-positive or V5-positive signal per DRG. Data are presented as mean ± SEM, analysed by unpaired t-test, **p < 0.01, ****p < 0.0001, n = 8 DRGs/group, with three sections evaluated per DRG to obtain the final value. (**E, F**) Immunoblotting results for GFP, PKA C-α, β-actin, and vinculin in DRG samples from animals injected with GFP only and those co-injected with db-cAMP 13 weeks post-injection. Values were normalised to vinculin. Two housekeeping proteins were utilised to enhance the reliability and accuracy of the results. Results are presented as mean density values ± SEM; n = 3 DRGs, with western blots repeated four times in separate experiments; ns > 0.5, *p < 0.05, **p < 0.01, analysed by unpaired t-test. (**G**) Heat maps illustrate Log_2_ fold changes (as Log_2_(2^-ΔΔCt)) for the expression of *Creb1* and *Itg*α*9*, normalised to the uninjured, uninjected control. Log_2_(2^-ΔΔCt) values were determined through qRT-PCR analysis. Results are presented as a heat map.

However, we observed a significant decrease in GFP and V5 signal intensity when AAVs were injected together with db-cAMP in DRG sections, as measured and analysed in ImageJ (Figure 4D). To further explore the hypothesis that db-cAMP might reduce the expression of the gene product carried by the AAV vector, we performed Western blot analyses on DRGs from animals in the GFP control and GFP+db-cAMP groups (Figure 4 E). In addition to detecting GFP protein, we measured PKA C-α, which is directly activated by db-cAMP, to confirm the presence of db-cAMP in the DRGs. The Western blot results showed an increased level of PKA C-α in the db-cAMP group, indicating the activity of db-cAMP within the DRGs. Interestingly, PKA C-α levels remained elevated in the DRGs even after the effects of db-cAMP had dissipated, suggesting that the increase in PKA C-α was a persistent effect independent of the transient db-cAMP activity.

A surprising finding was the sustained effect of PKA in DRG neurons, observed even 13 weeks after the co-application of both the AAV vector(s) and db-cAMP, as indicated by the elevated levels of PKA C-α. Notably, in the db-cAMP group, the GFP signal displayed a split band, suggesting minimal phosphorylation of GFP by PKA. Due to this split band, we could not determine whether GFP expression was downregulated or if the staining was influenced by phosphorylation. Further experiments will be required to clarify this mechanism.

Knowing that one of the functions of PKA is to regulate gene transcription^31^, we decided to investigate the expression of integrin α9 at different time points following the application of db-cAMP, both with and without the AAV vector. We chose the *Itga9* gene because of its crucial role in axon regeneration within inhibitory environments and the high availability of TaqMan probes for it, compared to the limited options available for GFP TaqMan probes. Using qRT-PCR, we aimed to confirm the long-term effects of db-cAMP observed through western blot analysis by examining the mRNA levels of *Creb1*, protein binding the cAMP response element, a DNA nucleotide sequence present in many viral and cellular promoters. The binding of CREB1 stimulates transcription.

Our qRT-PCR results revealed elevated levels of *Creb1* mRNA 42 days after application in both the GFP+db-cAMP injected group and the group receiving α9+K1in combination with db-cAMP, with no significant difference between the two treatment groups. In contrast, *Creb1* mRNA levels were significantly higher in the db-cAMP treated groups compared to those receiving only AAV-hSYN-α9-V5, where only physiological levels were expected. Additionally, we observed a significantly lower level of *Itg*α*9* mRNA when db-cAMP was co-injected with the AAV vector. This suggests that db-cAMP may function as a negative regulator of AAV vector transcription (Figure 4 G).

### Db-cAMP reduces the synthesis of TnC by neurons and astrocytes

From previous studies, we know that full integrin α9 activation relies not only on kindlin-1, but also on neuronal TnC^32^, and additionally astrocytic TnC is important for astrocytes to maintain the ramified presumably supportive regulatory astrocytic phenotype in scar tissue in the ischemic brain^21^. To further investigate whether application of db-cAMP can regulate the TnC production we first focused on the neuronal production of TnC. Using immunohistochemistry on the primary mouse DRG cell cultures and on rat DRG sections, we measured the relative intensity of TnC staining. The altered TnC expression pattern was conformed using western blots from DRG (13 weeks post application Our findings suggest that db-cAMP can downregulate the TnC production in neurons (Figures 5 A, B; Figure 6 A, B). This may explain why there was no synergy between the treatments *in vivo* in spite of the positive effects shown *in vitro* in Figure 2. This also explains why a synergistic effect was observed in vitro when cells were plated on TnC, as α9 activation depends on exogenous TNC. In contrast, this effect was absent when cells were plated on PDL, because α9 activation depends on neuron-produced TNC, which is downregulated by db-AMP.

**Figure 5.**
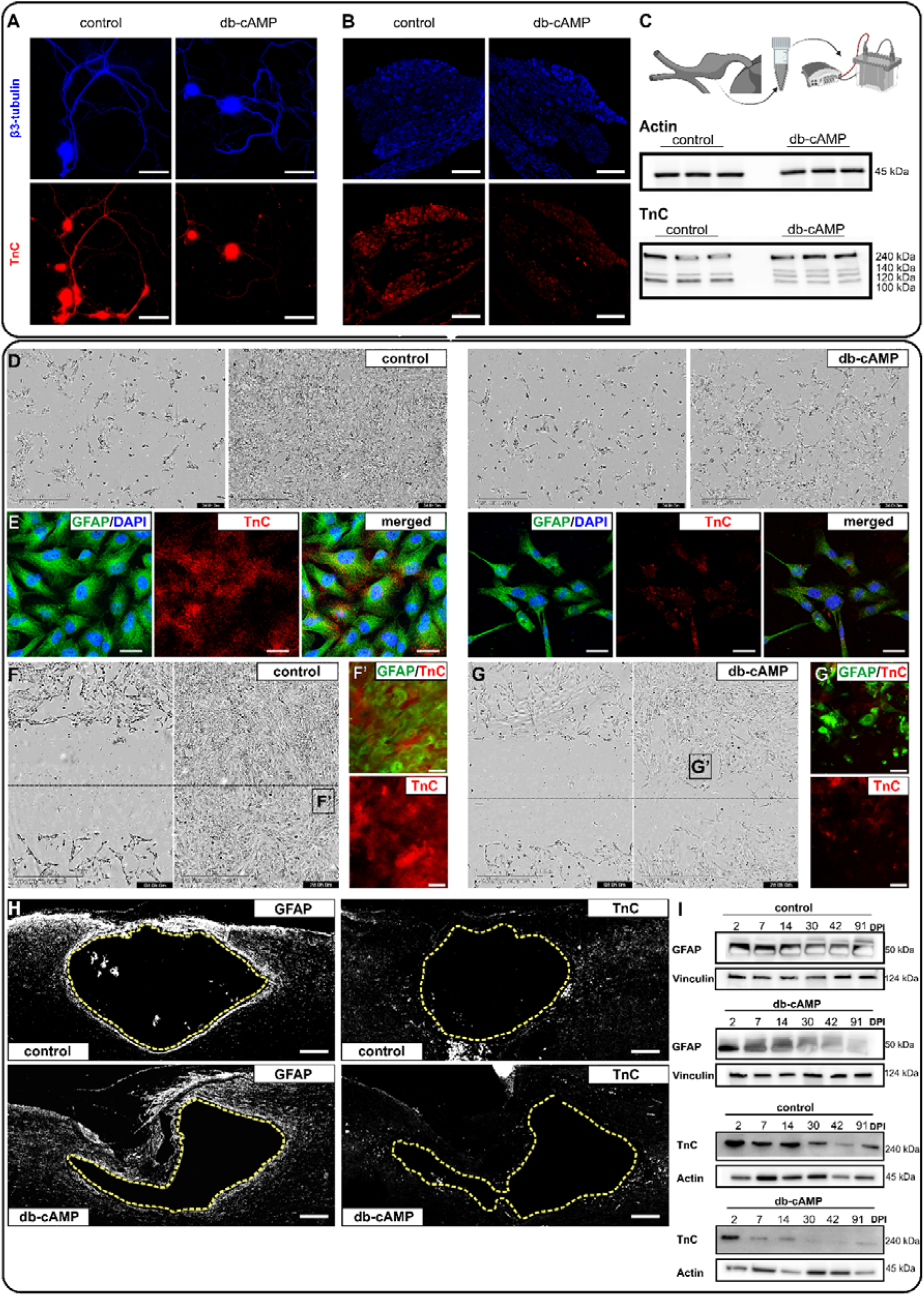
Db-cAMP reduces TnC synthesis in neurons and astrocytes. **(A**) Representative images of TnC in cultured primary mouse DRG neurons with or without db-cAMP treatment reveal reduced TnC signal in treated samples, while β3-tubulin remains unchanged. Scale bar: 70 µm. (**B**) Confocal images of DRG sections immunostained for TnC and β3-tubulin further confirm reduced TnC signal under db-cAMP treatment. Imaging parameters were consistent across samples. Scale bar: 400 µm. (**C**) Schematic and immunoblotting results show db-cAMP reduces TnC levels in DRGs. Western blot analyses reveal distinct TnC forms between treated and untreated groups, with β-actin serving as a control. (**D**) Incucyte images show altered astrocytic behaviour with db-cAMP treatment over 2 days. Treated astrocytes appear more fibrous and less adherent compared to control astrocytes. Scale bar: 400 µm. (**E**) Immunostaining of astrocytes post-cultivation shows a shift in TnC localization. Control astrocytes produce and secrete TnC, mostly at cell-cell contacts. Db-cAMP-treated astrocytes produce some TnC but fail to secrete it, with TnC retained in the cytoplasm. Scale bar: 50 µm. (**F**, **G**) Live imaging during scratch assays shows impaired migration and wound closure in db-cAMP-treated astrocytes compared to controls. (G’, F’) Immunostaining post-scratch confirms morphological changes and reduced TnC secretion. Scale bar: 600 µm (scratch assay), 50 µm (immunostaining). (**H**) Immunostaining of spinal sections 13 weeks post-injury shows no significant differences in TnC and GFAP expression between treated and untreated groups. Scale bar: 200 µm. (**I**) Immunoblotting of spinal cord samples from treated and control animals reveals consistent GFAP and TnC levels 13 weeks post-treatment, normalised to vinculin or β-actin based on protein kDa. These results show that db-cAMP reduces the production of TnC in neurons and astrocytes and alters the behaviour and migration of these cells in vitro and in vivo.

**Figure 6.**
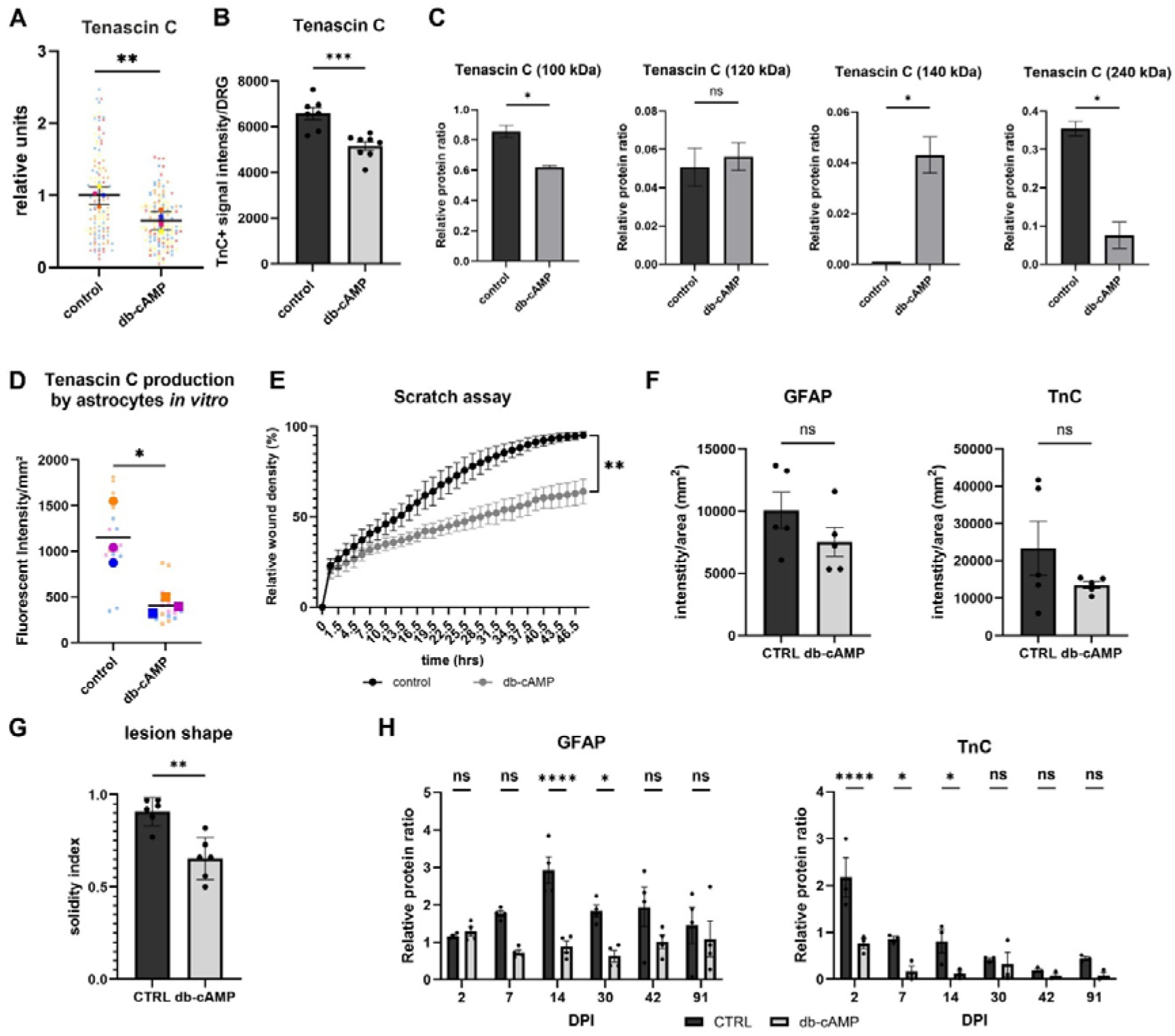
Db-cAMP reduces TnC synthesis by astrocytes and neurons. **(A)** Quantification of Figure 5A. The superplot shows relative signal intensity to assess db-cAMP’s effect on neuronal TnC synthesis in vitro. Individual data points (from three biological replicates) are colour-coded, with mean ± SEM overlaid. Significant reduction in TnC synthesis was observed (**p < 0.01, unpaired t-test). **(B)** Quantification of Figure 5B. Bar graph represents mean TnC intensity per DRG, with individual data points showing values from three sections averaged per DRG. Data are presented as mean ± SEM; ***p < 0.001, unpaired t-test. **(C)** Quantification of Figure 5C. Bar graph shows mean density values ± SEM for TnC from western blots. Results from three DRGs across four experiments indicate a significant reduction (*p < 0.05, unpaired t-test; ns > 0.5). **(D)** Quantification of Figure 5E. Superplot represents TnC signal intensity per cell coverage area in astrocytes treated with db-cAMP. Individual replicates are colour-coded, with mean ± SEM overlaid. Significant reductions were noted (*p < 0.05, unpaired t-test). **(E)** Quantification of Figure 5F. Graph shows relative wound density (%) over time from the scratch assay. Db-cAMP significantly reduced wound closure efficiency (**p < 0.01, two-way ANOVA, Šídák’s test; n = 3 experiments). **(F)** Quantification of Figure 5H. Bar graph shows TnC and GFAP intensity in spinal sections 13 weeks post-injury. Mean ± SEM values for five spinal cords (averaged from three sections each) show no significant differences (ns > 0.5, unpaired t-test). **(G)** The solidity index, quantifying lesion shape changes with db-cAMP treatment, shows significant differences (**p < 0.01, unpaired t-test; n = 6 spinal cords). **(H)** Quantification of Figure 5I. Bar graphs represent relative protein levels of TnC and GFAP normalised to vinculin or β-actin across time points. Db-cAMP treatment caused a significant reduction in TnC levels at certain time points (*p < 0.05, ****p < 0.0001, two-way ANOVA, Šídák’s test; n = 4 experiments).

Lastly, we examined the expression of TnC produced by astrocytes. Given that TnC plays a key role in wound healing and tissue remodelling following injury^21^ and noting the distinct differences in lesion shape seen in Figure 3 H and in Figure 5 H, I, we explored whether administering db-cAMP into the DRG might influence lesion formation.

To investigate this, we performed immunostaining for TnC and GFAP in spinal sections. No significant changes in the signal intensity of GFAP or TnC were observed 13 weeks after DRG injections. However, recognising that scar formation after SCI is a dynamic, time-dependent process of continuous remodelling^33^, we conducted western blot analyses of spinal cord tissue from the injury site, focusing on GFAP and TnC protein levels (Figure 5 I, Figure 6 H).

The western blot results revealed notable changes in both TnC and GFAP expression levels. TnC showed significant alterations at early post-injury stages (2-, 7-, and 14-days post-injury, DPI), while decreased GFAP levels were observed at intermediate timepoints, particularly at 14 and 30 DPI in the db-cAMP group (Figure 5 I, Figure 6 H). These findings suggest that the altered, irregular lesion shape may result from modified ECM expression in scar-forming cells following db-cAMP injections into the DRG (Figure 6 G).

To determine whether the db-cAMP-mediated effects on astrogliosis represent a conserved mechanism, we studied its effects on a cultured human astrocytic cell line. First, we treated astrocytes with and without db-cAMP, focusing on TnC production after 2 days. Treatment with db-cAMP led to a significant decrease in TnC production by astrocytes, along with altered localisation. In the absence of db-cAMP, astrocytes secreted TnC, but with db-cAMP treatment, TnC remained detectable in the cytoplasm by immunostaining and was not secreted (Figure 5 D, E; Figure 6 D). In addition, db-cAMP treatment affected astrocytic proliferation and morphology, as astrocytes did not adopt the typical tile-like shape when in contact with each other under db-cAMP conditions (Figure 5 D, E). Our preliminary data (see Supplementary Material, Fig. S1) also show similar results in primary mouse astrocytic cultures, suggesting that the effect on astrocytic TnC production appears to be evolutionarily conserved in the mammalian CNS. These observations are consistent with previous findings that TnC is essential for maintaining astrocyte proliferation and morphology in primary cultures^34^.

We then performed a scratch assay using Incucyte to assess cell migration. After 2 days, untreated astrocytes had migrated into the scratch area and completely closed the wound with no visible remnants. In contrast, db-cAMP-treated astrocytes failed to migrate and close the *in vitro* wound within the same time frame. These results suggest that the effect of db-cAMP on TnC expression and astrocyte behaviour may be conserved throughout the mammalian CNS (Figures 5 F, G; Figure 6 E).

## Discussion

In this study, we present a combined strategy for axon regeneration after thoracic SCI in rats. Our approach combines db-cAMP-mediated conditioning with AAV-mediated overexpression of activated α9 integrin to promote axonal growth and repair. Traditionally, axon regeneration after SCI has relied on the classical conditioning injury paradigm, which has been considered the gold standard for sensory axon regeneration since its discovery decades ago^10,35,36^. This paradigm facilitates the expression of a regeneration-associated gene (RAG) programme, which is essential for the initiation of neuronal repair and regeneration. Without activation of the RAG programme, significant regeneration is unlikely. Preconditioning, such as db-cAMP treatment, has been shown to prime sensory neurons for regeneration by inducing the expression of this RAG programme, resulting in limited local regeneration at the site of injury. The molecular changes associated with α9 integrin-induced regeneration, including mRNA changes, are discussed in detail in Cheah et al.^18^.

We found that the combination of db-cAMP, which mimics a conditioning lesion^13^, with overexpression of activated α9 integrin enhanced the regenerative potential of DRG neurons, resulting in increased neurite outgrowth in vitro. Specifically, our in vitro studies showed that the combination of AAV-mediated overexpression of integrin α9 with db-cAMP administration significantly increased neurite length on a mixture of TnC and Agg, indicating enhanced axonal growth and regeneration potential. However, this promising result did not translate into equivalent in vivo efficacy when both AAV integrin α9 and db-cAMP were injected into DRGs after SCI. Several factors may contribute to this discrepancy.

In a controlled *in vitro* environment, neuronal growth conditions are optimised, such as uniform substrate attachment and a stable environment, which are not present in the complex *in vivo* environment. An important factor that may have contributed to the observed differences is the exogenous addition of TnC in the *in vitro* setup, which is likely to have masked the regulatory effects of db-cAMP on TnC expression and production at the cellular level. In addition, db-cAMP was present throughout the *in vitro* experiments, whereas in our SCI experiments db-cAMP was injected only at the time of injury. In addition, the *in vivo* environment is dominated by inhibitory factors such as chondroitin sulphate proteoglycans, inflammatory responses and scarring, all of which may impede axonal regeneration despite the promising *in vitro* results^37^. Finally, effective delivery of AAV integrin α9 and maintenance of its expression in the hostile post-injury environment are significant challenges that may limit the expected regenerative outcomes. In previous studies, only about 50% of axons successfully crossed the lesion site^16^, highlighting the need for more refined therapeutic strategies to overcome these biological barriers *in vivo*.

This study proposes a potential mechanism to explain why the combination of AAV integrin α9 and db-cAMP was less effective *in vivo*. First, the differences in signal intensity observed in the DRGs indicated lower production of GFP and/or V5. One hypothesis is that increased PKA activity due to elevated cAMP levels may have phosphorylated both GFP and V5. These proteins contain serine, threonine and tyrosine residues that are potential phosphorylation sites for kinases. However, studies suggest that GFP is minimally phosphorylated by PKA, a serine/threonine kinase, suggesting that phosphorylation is unlikely to have significantly affected the experiment. For PKA to phosphorylate a target, the surrounding sequence must align with its recognition motif (RRXS/T)^38^. In the V5 tag sequence, neither serine nor threonine is part of a strong PKA recognition motif, making it unlikely that PKA-mediated phosphorylation would interfere with V5 detection during immunostaining. Similarly, phosphorylation is unlikely to affect GFP fluorescence as the chromophore responsible for fluorescence is well protected within the protein structure^39^. Unless phosphorylation induces a major structural change, fluorescence should remain unaffected. Although the western blot revealed a double band in GFP, likely due to the db-cAMP-induced increase in PKA levels in the DRGs, it remains unclear whether the reduced fluorescence signal in both V5 and GFP following db-cAMP treatment is due to minimal phosphorylation - potentially affecting detection - or a decrease in GFP and V5 expression mediated by db-cAMP. Further investigation is required to clarify these potential effects.

RT-qPCR analysis revealed reduced levels of *Itga9* mRNA when AAV-hSYN-α9 was co-administered with db-cAMP. To our knowledge, there are no studies showing the effect of elevated PKA levels on AAV1, however, phosphorylation on AAV2 has previously been shown to negatively affect intracellular trafficking and transduction efficiency of recombinant AAV2 vectors^40^. Phosphorylated AAV vectors show a significant reduction in transduction efficiency—approximately 68% for single-stranded AAV (ssAAV) and 74% for self-complementary AAV (scAAV)—despite entering cells as efficiently as non-phosphorylated vectors^40^. This decreased transduction is attributed to reduced intracellular trafficking from the cytoplasm to the nucleus, which is associated with increased ubiquitination and subsequent degradation of the capsids^40^.

Despite a single injection of db-cAMP, we observed a long-term effect of db-cAMP on both sensory neurons and the CNS injury site. The prolonged increase in PKA activity observed even 13 weeks after db-cAMP injection in DRG neurons suggests significant long-term effects on neuronal signalling. Although db-cAMP acts as a stable cAMP analogue, it is unlikely that it would have persisted in the DRGs for 13 weeks. Instead, the sustained increase in PKA activity is likely to be due to cAMP-independent mechanisms, where changes in PKA regulatory subunits or feedback loops maintain the activation of the kinase^41,42^. This phenomenon has been observed in several contexts, including long-term synaptic plasticity^42^, where PKA can remain active without the need for continuous cAMP presence, and is likely to contribute to the prolonged effects observed in our study.

Our data show that db-cAMP application led to a significant reduction in TnC levels in both neurons and astrocytes. While TnC is primarily produced by astrocytes^43^, fibroblasts and other non-neuronal cells, neurons can express it during development or in response to injury^23^. Neuronal TnC supports axonal growth, cell migration and synaptic plasticity, and its expression is typically low in adults, but increases after injury to play a critical role in tissue remodelling, neurogenesis, synaptic modulation and CNS repair^20,24,44–46^. After CNS injury, TnC is upregulated, mainly by astrocytes, in response to various stimuli, although its expression is spatially and temporally restricted^21,44^. TnC regulates gene expression and cell migration through integrin interactions and other signalling pathways, and CREB has been identified as a potential regulator of TnC expression^47^. CREB can act as both a silencer and an enhancer of gene expression^48^, which may explain its silencing effect on TnC when overexpressed by db-cAMP. In vitro studies in neurons and astrocytes confirmed the direct involvement of the db-cAMP/PKA/CREB pathway in modulating TnC expression.

Notably, db-cAMP had a significant effect on astrocytes, influencing their proliferation and migration in wound healing assays, reflecting the close relationship between TnC and astrocytic migration^34^. TnC is essential for maintaining the proliferation and correct morphology of astrocytes in primary culture and plays a key role in limiting excessive gliosis while maintaining the supportive phenotype of astrocytes in the ischaemic brain^21,34^.

In vivo, we observed reduced GFAP reactivity and a smaller cavity at the SCI site in db-cAMP-treated animals, supporting previous findings of scar reduction. Interestingly, the effects on GFAP at early time points post-SCI and the influence on astrocytic TnC at the lesion border were observed even though db-cAMP was applied only to the DRGs. The mechanisms underlying this localised, yet widespread effect remain unclear, although it is possible that the conditioning effect of db-cAMP may have affected the lesion border itself. Previous conditioning lesions have been shown to alter the inflammatory response, neuroplasticity and vascular changes, all of which could contribute to changes at the lesion border and surrounding tissue. These effects, induced by previous injury or conditioning, may lead to changes in the lesion environment, thereby influencing the characteristics of the injury and tissue regeneration at the border^49^. Further investigation is required to fully elucidate the mechanisms involved.

Collectively, these data suggest the importance of the presence of TnC, both endogenously synthesised by neurons and exogenously produced by astrocytes, for the proper astrocytic barrier around the lesion and for α9-mediated regeneration.

## Conclusion

In this study, we successfully replicated previously observed results and demonstrated that activated integrin α9 enhances neurite outgrowth *in vitro,* and that AAV-mediated expression of activated α9 promotes the sensory pathway reconstruction *in vivo*. In addition, we confirmed the beneficial effect of db-cAMP on axon growth following spinal cord injury (SCI) in rodent models. However, the study also showed that combining db-cAMP with activated integrin α9 did not enhance sensory axon regeneration. This finding is likely due to two key factors: (i) the db-cAMP/PKA/CREB pathway may reduce AAV-mediated gene expression, and (ii) db-cAMP downregulates TnC expression both *in vitro* and *in vivo*, in neurons and astrocytes. This inhibition limits extensive regeneration of sensory axons and prevents the formation of long-range, topographically correct connections within the spinal cord. Overall, this study uncovers a previously unknown effect of the cAMP/PKA/CREB pathway in regulating TnC expression and highlights its potential role in influencing axonal regeneration.

## Materials and methods

### Preparation of AAV vectors

To produce the virus, HEK293T cells were transfected with individual plasmids and helper genes^50^. The cells were cultured for a period of three days. Then the cells were lysed. This was done with three freeze and thaw cycles. After centrifugation, the crude supernatant was ultracentrifuged at 490000 x g, 16°C for 70 min through iodine hexanol gradients (15%, 25%, 40% and 60%) using a Beckman 70Ti rotor. An Amicon Ultra-15 (Millipore) was then used to collect and concentrate the virus. Virus titre was then determined by quantitative real-time PCR at levels of 2.34×10^12 GC/ml for AAV1-α9-V5 and 4.99×10^12 GC/ml for AAV1-Kindlin1-GFP. AAV1-SYN-GFP (3×10^13 GC/ml) is supplied by Vigene, distributed by Charles River (CV17001-AV1).

A key difference between in vitro and in vivo experiments lies in the duration of observation. In vitro studies capture immediate cellular responses, while in vivo experiments extend over a longer period to assess the sustained regenerative effects of treatments. This extended timeframe is critical for understanding the long-term impact of therapies on neuronal recovery after SCI. Although in vitro studies offer valuable insights into initial mechanisms, long-term in vivo observations are essential to evaluate the true efficacy of therapeutic strategies.

### Cell cultures and experimental animals

*In vitro* experiments were initially performed in primary mouse cultures due to their availability and ease of colony maintenance. Subsequent *in vivo* experiments were performed in rats, as their larger anatomical size facilitates precise injection into the DRG and allows for more reliable behavioural assessment. Finally, experiments with astrocytes used a human immortalised cell line to provide translational relevance.

### Cell culture of DRG neurons/cortical astrocytes

Adult 3-month-old WT C57BL/6 mice were overdosed with isoflurane and decapitated. All DRGs were then removed and dissociated in 0.2% collagenase (Sigma-Aldrich; # C9407) for 2 hours and 0.1% trypsin (Sigma-Aldrich; #T0303) for 10 minutes in DMEM (Thermo Fisher Scientific; #10566016) at 37°C. The cells were subjected to mechanical trituration and centrifugation on a 15% BSA (Sigma-Aldrich; #A9418) gradient. The cells were plated in the following culture medium: 1% ITS (Sigma-Aldrich; #I2521), 1% PSF (Sigma-Aldrich; #P4333), 10 ng/ml NGF (Sigma-Aldrich; #N8133), and 0.5 µg/ml mitomycin-C (Sigma-Aldrich; #M4287) in DMEM on coated glass coverslips at a density of 1 × 10^4 cells/condition. The coverslips were treated with either 20 µg/ml PDL (Sigma-Aldrich; #P1149) or 1 µg/ml PDL + 10 µg/ml TnC (Sigma-Aldrich; *#*CC115) + 10 µg/ml aggrecan (Agg) (Sigma-Aldrich; #A1960). Under the different conditions, cells were either left untreated or treated with 2 mM db-cAMP (Sigma-Aldrich; #D0260) for 5 days in vitro (DIV). Cortical astrocytes were generously provided by Eickholt’s lab (Charité, Berlin, DE), and a single experiment was performed to generate the preliminary data supporting the results presented.

### Cell culture transduction of DRG neurons

Dissociated DRG neurons were incubated with 1 × 10^9 TU/ml AAV-SYN-GFP, AAV-SYN-α9-V5 or AAV-SYN-α9-V5 + AAV-CMV-K1-GFP for 72 hours at 37°C to induce viral transduction. To ensure protein expression, the medium was then changed and the cells incubated for a further 48 hours.

### Astrocyte cell culture, proliferation and scratch wound assays

Astrocyte experiments were performed using immortalised human astrocytes (IM-HA P10251-IM, Innoprot, Spain). Cells were cultured in T75 flasks (Nunc, Thermo Fisher Scientific) coated with poly-L-lysine (Sigma-Aldrich; # P4707) in a CO2 incubator (MCO-170AICUVH-PE, Panasonic, Osaka, Japan) at 37°C. IM-HA cells were cultured in a medium consisting of DMEM (Thermo Fisher Scientific; #D0697) supplemented with 10% heat-inactivated foetal bovine serum (Thermo Fisher Scientific; #16140071), 1% non-essential amino acids (Thermo Fisher Scientific; #11140050), 1% N-2 supplement B (Stemcell Technologies; #7156), 1% L-glutamine (Sigma-Aldrich; #G7513), 1% penicillin-streptomycin mixture (Thermo Fisher Scientific; #15140122), and 0.5% Normocin (InvivoGen). Cells were passaged at 80% confluence with trypsin/EDTA solution (Thermo Fisher Scientific; #R001100).

For the proliferation assay, 24-well plates (TPP, Trasadingen, Switzerland) with coverslips were coated with poly-L-lysine. One million cells were seeded per 24-well plate. One day after seeding, the medium was replaced in all wells (n = 12 per experiment, repeated in three independent experiments). The treated wells (n = 6) were further supplemented with 2 mM db-cAMP (Sigma-Aldrich, St. Louis, MO, USA). Plates were placed in an incubator (Sartorius, Göttingen, Germany) and cell confluence was measured every hour for the next 2 days.

For the scratch assay, 96-well plates (TPP, Trasadingen, Switzerland) were coated with poly-L-lysine. One million four hundred thousand cells were seeded per 96-well plate. One day after seeding, the medium was changed in all wells (n = 40 per experiment, repeated in three independent experiments) and the treated wells (n = 20) were supplemented with 2 mM db-cAMP. After the medium change, a scratch wound was made using standardised equipment (Sartorius, Göttingen, Germany). The 96-well plates were then placed in an Incucyte (S3, Sartorius, Göttingen, Germany), and wound closure was measured every hour for 2 days. Both the proliferation and scratch assays were repeated three times for statistical analysis.

### Experimental animals and animal surgeries

Sixty-seven female Lister Hooded rats (150-175 g; Envigo) were used in this study. Rats were housed in groups of three in cages with a 12-hour light-dark cycle under standard conditions of temperature (22 ± 2°C) and humidity (50% ± 5%). Only female rats were used in this study to reduce the potential negative effects of rapid weight gain, as observed in male rats in long-term studies, particularly during housing and behavioural testing. Rats had free access to tap water and food ad libitum. All procedures were approved by the Ethics Committee of the Institute of Experimental Medicine of the ASCR and were performed in accordance with Act No. 77/2004 of the Czech Republic (ethics approval number: AVCR 4386/2022 SOV II). Based on previous studies, the number of animals for each experiment was statistically optimised to achieve its reduction in accordance with European Commission Directive 2010/63/EU, and every effort was made to minimise pain and suffering. Each animal was numbered and randomly assigned to either the control or experimental group.

The sample sizes in this study were determined based on a power analysis targeting a large effect size (f = 0.4) with an α level of 0.05 and 80% power, suggesting an optimal range of 6–12 animals per group for behavioural tests. We included the following groups: GFP alone (GFP control group), db-cAMP with GFP (GFP+db-cAMP group) (n = 12 each) and α9-K1 alone (α9-K1 group) and a9-K1 with db-cAMP (a9-K1+db-cAMP) (n = 6 each). In the subsequent time-course study to further validate our findings, a sample size of 3 animals per time point was used, consistent with the 3Rs principles.

To maintain blinding, all surgeries were performed by two independent investigators, one performing the DRG injection and the other inducing the SCI. Surgery was performed under inhalational anaesthesia with isoflurane (1.8-2.2%; Baxter Healthcare Pty Ltd; #26675-46-7) in 0.3 l/min oxygen and 0.6 l/min air. All animals received buprenorphine (Vetergesic® Multidose) subcutaneously at a dose of 0.2 mg/kg body weight. Using sterile surgical equipment, two DRGs (L4, L5) were exposed on the left side and 1.5 μL of viral vector at a working titre of 2×1012 GC/ml alone or mixed with 10mM db-cAMP (Sigma-Aldrich; #D0260) was manually injected using a Hamilton syringe with custom-made needles (Hamilton; specification: 33 gauge, 12mm, PST3). Simultaneously, a laminectomy was performed at T10. Small slits were made in the dura using a U100 insulin syringe (B Braun Omnican 50 insulin syringe; #9151117S), and the dorsal columns were crushed using fine Bonn forceps (Fine Science Tools). The tips of the Bonner forceps were held on either side of the dorsal columns where the small slits had been made, pressed 1 mm deep into the spinal cord and then held firmly together for 15 seconds. Many previous studies have shown that this injury results in a complete transection of the dorsal columns down to the central canal^51^. Immediately after the injury, the muscles and skin were sutured. At the end of the experiment, the animals were first given a lethal dose of ketamine and xylazine and then either perfused or decapitated. Their tissues were then processed according to the specific requirements of the subsequent experiments.

Animals were anaesthetised intraperitoneally with a lethal dose of a combination of ketamine (100 mg/kg) and xylazine (20 mg/kg) and perfused intracardially with 4% paraformaldehyde (PFA) in 1X-PBS and the tissue used for immunohistochemistry. Animals whose tissues were used for western blot and qPCR analyses were anaesthetised intraperitoneally with a lethal dose of ketamine (100 mg/kg) and xylazine (20 mg/kg), decapitated, and the DRGs and spinal cord quickly collected on dry ice. The tissues were then stored at −80°C until processed. One uninjured and untreated animal was sacrificed to provide a negative control for western blot analysis.

### Triple immunosuppressive therapy

Triple drug immunosuppression was used to prevent potential AAV-mediated DRG toxicity^52^. All animals were treated intraperitoneally with cyclosporine A (10 mg/kg) (Novartis, East Hanover, NJ, USA), orally with azathioprine sodium (4 mg/kg) (Aspen Pharma Trading Ltd., Dublin, Ireland) and intramuscularly with methylprednisolone (2 mg/kg) (Pfizer Inc., New York, Ny, USA). Immunosuppressive therapy was started 1 week before surgery. The immunosuppressants were administered daily for the first month, then every other day for one month, and finally reduced to twice weekly.

### Behavioural testing

The von Frey test, the Hargreaves test and the tape removal test were performed every two weeks for 12 weeks to ensure that the animals did not learn the task independently of the sensory reflexes. Basso, Beattie and Bresnahan (BBB) test was performed weekly to regularly check motor function and consistency of the lesion size. Animals were brought into the testing room at least 20 minutes prior to testing to allow them to become accustomed to the environment. Each rat was measured 3 times prior to surgery to provide a baseline for each test. Behavioural testing was performed by two independent investigators.

### Von Frey test

Touch sensitivity was measured using the electronic von Frey test (IITC Inc., Life Science Instruments, Woodland Hills, CA, USA). Animals were placed in the Plexiglas enclosure on a gridded floor stand (IITC Inc., Life Science Instruments, Woodland Hills, CA, USA) for at least 20 min prior to testing to allow for acclimation. The left and right hind paws were then stimulated by a slowly rising probe with a rigid plastic tip touching the centre of the hind paw. The pressure was increased until the nociceptive response occurred in the form of withdrawal of the paw. The value was then recorded. Each hind paw was measured 5 times. The lowest and highest of these five values were discarded and the remaining three values were averaged.

### Hargreaves test (Plantar test)

SCI-mediated changes in heat sensation were measured using the Ugo Basile plantar heat test apparatus (Comerio VA, Italy). Rats were placed in a Plexiglas box with a fibreglass bottom and acclimatised for 30 min. An infrared radiator (Ugo Basile) was then placed directly under the hind paw pad. The time at which the rat withdrew its paw was recorded. The timer was automatically activated when the stimulus was applied. The infrared stimulus was automatically turned off after 30 seconds to avoid injury to the animal. Five trials were performed on the left and right hind paw, with a minimum of 3 min rest between each trial. The average withdrawal time was calculated by taking the average of the three trials after deleting the lowest and highest values for each animal.

### Tape removal test

For the tape removal test, each rat was trained in 5 sessions and then tested every other week after surgery. The rat was then removed from its home cage, placed in an empty test cage and allowed to acclimate for 15 minutes. A small piece of tape (approximately 1 square centimetre) was placed on the paw. Three trials were performed on each hind paw with a minimum of 3 minutes rest between trials. The experimental and control paws were tested simultaneously. The time at which the animal first noticed the tape was recorded. If the rat did not notice the tape within 5 min, the tape was removed and a time of 5 min was recorded. Paper tape (TimeMed Labelling Systems, Inc.; Fisher Scientific; #NC9972972) was used for this test.

### Basso, Beattie and Bresnahan (BBB) test

The Basso, Beattie and Bresnahan (BBB) open field test was used to assess locomotor performance in rats. Consistency of lesion size between animals was the primary objective of this test. Rats were placed in an open field arena. The open field arena was surrounded by a rectangular pen. Results were scored on a range of 0-21 points; from complete lack of locomotor ability (0) to healthy rat-like locomotor ability (21)^53^.

### Immunostaining

#### Immunocytochemistry

After 5DIV, cells were fixed with 4% paraformaldehyde (PFA) for 15 minutes, followed by incubation with immunoblocking solution consisting of 1% BSA in PBS and 2% Triton for 2 hours. The primary antibodies used in this study are listed in Table 1. The primary antibodies were kept overnight at 4°C in PBS with 1% BSA and 2% Triton. The next day, the cells were washed and incubated with Alexa Fluor®-conjugated secondary antibodies for 2 hours to detect the primary antibodies. The goat anti-host antibody of each primary antibody was conjugated with Alexa Fluor® 405, 488, 546 and 594 (1:400; 2 h; room temperature (RT)). FluorSave™ (Calbiochem; #345789) was used to mount the coverslips.

**Table 1.** Primary antibodies and labelling agents used in this study, detailing their respective epitopes, host species, providers, catalogue and LOT numbers, as well as application-specific dilutions.

| 1° Antibody/labelling agent | Epitope | Host | Company | LOT | Dilution/incubation |
| --- | --- | --- | --- | --- | --- |
| Anti-V5 | V5 tag on the α9 integrin | Mouse monoclonal | Invitrogen #R960-25 | 2901472 | ICC: 1:500; O/N<br>IHC: 1:500; 48h |
| Anti-GFP | Green fluorescent protein (GFP) | Chicken polyclonal | Invitrogen #A10262 | 2738237 | IHC: 1:500; 48h<br>WB: 1:1000; O/N |
| Anti-GFP | Green fluorescent protein (GFP) | Rabbit polyclonal | Invitrogen #A-6455 | 2720383 | IHC: 1:500; 48h |
| Anti-β3 tubulin | β3 tubulin | Mouse monoclonal | Abcam #ab78078 | 1009304-10 | IHC: 1:800; 48h |
| Anti-β3 tubulin | β3 tubulin | Rabbit monoclonal | Abcam #ab68193 | 1024396-2 | ICC: 1:1,200; O/N<br>IHC: 1:800; 48h |
| Anti-Tenascin C | Tenascin C (KAF12) | Rabbit polyclonal | courtesy of the Faissner lab<sup>22</sup> | N/A | ICC: 1:400; O/N |
| Anti-Tenascin C | Tenascin C | Rabbit monoclonal | Cell Signaling #33352 | N/A | ICC: not used<br>IHC: 1:600; 48h<br>WB: 1:1,000; 48h |
| Anti-GFAP | Glial fibrillary acidic protein | Chicken polyclonal | Abcam #ab134436 | 1056567-3 | ICC: 1:500; O/N<br>IHC: 1:500; 48h |
| DAPI | 4',6-diamidino-2-phenylindole | N/A | Invitrogen #D1306 | 2274787 | ICC: 1:3,000; O/N<br>IHC: 1:5,000; 48h |
| Anti-β-Actin | β-Actin | Rabbit monoclonal | Cell Signaling #12620 | N/A | WB: 1:5,000; O/N |
| Anti-GFAP | Glial fibrillary acidic protein | Rabbit monoclonal | Cell Signaling #12389 | N/A | WB: 1:10,000; O/N |
| Anti-Vinculin | Vinculin | Rabbit monoclonal | Cell Signaling #13901 | N/A | **WB:** 1:1,000; O/N |
| anti-PKA C-α | Protein kinase A C-α | Rabbit polyclonal | Cell Signaling #4782 | N/A | **WB:** 1:1,000; O/N |

| 1° Antibody/labelling agent | Epitope | Host | Company | LOT | Dilution/incubation |
| --- | --- | --- | --- | --- | --- |
| Anti-V5 | V5 tag on the $\alpha 9$ integrin | Mouse monoclonal | Invitrogen #R960-25 | 2901472 | ICC: 1:500; O/N<br>IHC: 1:500; 48h |
| Anti-GFP | Green fluorescent protein (GFP) | Chicken polyclonal | Invitrogen #A10262 | 2738237 | IHC: 1:500; 48h<br>WB: 1:1000; O/N |
| Anti-GFP | Green fluorescent protein (GFP) | Rabbit polyclonal | Invitrogen #A-6455 | 2720383 | IHC: 1:500; 48h |
| Anti-$\beta 3$ tubulin | $\beta 3$ tubulin | Mouse monoclonal | Abcam #ab78078 | 1009304-10 | IHC: 1:800; 48h |
| Anti-$\beta 3$ tubulin | $\beta 3$ tubulin | Rabbit monoclonal | Abcam #ab68193 | 1024396-2 | ICC: 1:1,200; O/N<br>IHC: 1:800; 48h |
| Anti-Tenascin C | Tenascin C (KAF12) | Rabbit polyclonal | courtesy of the Faissner lab<sup>22</sup> | N/A | ICC: 1:400; O/N |
| Anti-Tenascin C | Tenascin C | Rabbit monoclonal | Cell Signaling #33352 | N/A | ICC: not used<br>IHC: 1:600; 48h<br>WB: 1:1,000; 48h |
| Anti-GFAP | Glial fibrillary acidic protein | Chicken polyclonal | Abcam #ab134436 | 1056567-3 | ICC: 1:500; O/N<br>IHC: 1:500; 48h |
| DAPI | 4',6-diamidino-2-phenylindole | N/A | Invitrogen #D1306 | 2274787 | ICC: 1:3,000; O/N<br>IHC: 1:5,000; 48h |
| Anti-β-Actin | β-Actin | Rabbit monoclonal | Cell Signaling #12620 | N/A | WB: 1:5,000; O/N |
| Anti-GFAP | Glial fibrillary acidic protein | Rabbit monoclonal | Cell Signaling #12389 | N/A | WB: 1:10,000; O/N |
| Anti-Vinculin | Vinculin | Rabbit monoclonal | Cell Signaling #13901 | N/A | WB: 1:1,000; O/N |
| anti-PKA C-α | Protein kinase A C-α | Rabbit polyclonal | Cell Signaling #4782 | N/A | WB: 1:1,000; O/N |

#### Immunohistochemistry

PFA-fixed samples were cryoprotected by immersion in increasing concentrations of sucrose (10%, 20% and 30% in PBS). Tissue was then embedded in OCT mounting medium (VWR; #03820168). Sections were cut at 10 μm (DRG) and 20 μm (spinal cord) on positively charged slides (VWR; #630-1324) on a cryostat (Thermo Scientific, Cryostar NX70). The permeabilization step was performed in 0.5% Triton X-100 (Sigma; #T8787) for 2 hours at room temperature (RT) and the immunoblocking step was performed in 0.2% Triton X-100, 10% ChemiBLOCKER (Millipore; #2170) and 0.3M glycine (Sigma-Aldrich; #GE17-1323-01) for a further 2 hours at RT. After the immunoblocking step, sections were incubated with the following primary antibodies. The primary antibodies used in this study are listed in Table 1. For primary antibody detection, goat anti-host antibodies of the respective primary antibodies conjugated with Alexa Fluor 405, 488, 594 and 647 (1:300; 4 h, RT, Invitrogen) were used. Sections were then washed with 0.2% Triton X-100 in PBS and mounted with Mowiol mounting medium (Carl Roth; #0713.2). DABCO was added to reduce bleaching of the fluorescence signal (Carl Roth; #0718.1).

#### Histology

Luxol Fast Blue (LFB) staining was used to visualise the white and grey matter^54^ to compare lesion sizes in 3 randomly selected animals from each group, and to potentially exclude an animal if the lesion varied. No animal was excluded based on lesion size. Images were acquired using a LEICA CTR 6500 microscope with FAXS 4.2.6245.1020 software (TissueGnostics, Vienna, AT).

#### Tissue Clearing

Four randomly selected DRGs and three randomly selected spinal cord samples from each group were dehydrated through an ethanol (EtOH) dilution series (30%, 50%, 70%, 100%) with 2% Tween20 (Sigma-Aldrich; #P1379) and then delipidated in dichloromethane with EtOH (2:1). Samples were rehydrated with a series of decreasing EtOH concentrations (70%, 50%, 30%), always with 2% Tween20. A solution of 0.2% Triton in PBS, 0.3M glycine and 10% DMSO was then used to permeabilise the samples. Samples were blocked in 0.2% Triton in PBS, 0.3M glycine, 10% DMSO and 10% ChemiBLOCKER (2 days, 37°C). After blocking, DRG samples were incubated in the blocking solution with the following primary antibodies: anti-GFP (Invitrogen; #A-6455, 1:400, 2 days, 37°C) and anti-V5 (Invitrogen; #R96025, 1:400, 2 days, 37°C). Secondary antibodies (1:300) were prepared in blocking solution. These were used for 3 days at 37°C. Samples were cleared in ethyl cinnamate^55^ (Sigma-Aldrich; #112372) at RT (DRG for 2 hours; spinal cord for 24 hours). After clearing and imaging, samples were rehydrated in EtOH solutions of decreasing concentration (100%, 70%, 50%, 30%), washed in PBS and cryoprotected in sucrose solution (10%, 20%, 30%). Sections were cut and reused. UV light was used to remove the fluorescence signal. Re-use of samples reduced the number of animals required for this study.

#### Microscopy and image analysis

A Zeiss Axio Observer D1 (Zeiss, Oberkochen, Germany) inverted phase contrast fluorescence microscope was used for fluorescence imaging of DRG cell cultures at 40x magnification. Maximum neurite length was measured using ImageJ software^56^. Quantification of neurite outgrowth was performed from at least three different experiments. Comparisons were made between non-transduced (NT), GFP-transduced (GFP), α9-transduced (α9) and α9-kindlin1-transduced (α9K1) on permissive (PDL-coated coverslips) and inhibitory (TnC/Agg) media. Relative signal intensity was used to measure the effect of db-cAMP on neuronal TnC synthesis *in vitro*.

The regeneration index was analysed using an Axioskop 2 plus microscope (Zeiss, Oberkochen, Germany). The number of axons was counted at three random positions caudal and rostral to the lesion using an eyepiece grid by counting the number of axons crossing a line on the grid. The number rostral to the lesion was then divided by the number caudal to the lesion per section, and 5 sections per spinal cord were counted. The 5 values for an animal were then averaged.

A Zeiss LSM 880 (Zeiss, Oberkochen, Germany) confocal microscope was used to image DRG and spinal sections. Images were acquired under the same experimental conditions, including immunohistochemistry and microscopy settings (the same laser power, gain and airy units), for each staining set. The effect of db-cAMP on the scar was assessed by intensity measurement sampling - three random positions around the lesion were measured using Image J^56^, the three values were then averaged. All analyses were performed by two independent scientists. Values from both investigators were then pooled and the mean was calculated. Mean signal intensities were also measured in three randomly selected DRG sections, by two independent scientists. The cell counter plugin in Fiji was used to assess the number of transduced cells.

A Zeiss Lightsheet Z.1 (Zeiss, Oberkochen, Germany) was used for visualization of cleared DRG and spinal tissue. Huygens software (Scientific Volume Imaging, Hilversum, Netherlands) was used to correct chromatic aberration, which resulted in a shift between different channels in the Z-axis of the 3D image. Imaris software (Oxford Instruments, Abingdon, UK) was used for further analysis of the number of V5+/GFP+ cells in the DRGs and to generate 3D reconstructions of the spinal cords.

For all staining, the mean grey value of the corresponding secondary control sections was subtracted from each individual measurement to account for non-specific background staining. Identical settings and staining conditions were used for all images used for comparison. No image processing was performed. For publication resolution, only brightness and contrast were adjusted using Affinity V2 software. All raw data are available in the ASEP repository, Czech Academy of Sciences, in accordance with the principles of open science and scientific integrity.

#### Protein isolation and Western blot

The spinal cord and DRGs were rapidly removed on dry ice and stored at −80°C until protein isolation. Tissue segments around the injury site were cut into pieces and homogenised in RIPA Lysis and Extraction Buffer (Thermo Scientific™; #89900). The buffer contained the protease inhibitor cOmplete™ Protease Inhibitor Cocktail (Roche; #04693116001) and the phosphatase inhibitor PhosSTOP™ EASYpack (Roche; #05892970001). Samples were shaken on an orbital shaker for 30 minutes in a cold room (4°C). The samples were then vortex shaken and centrifuged at 15,000 × g for 20 minutes at 4 °C. The supernatant was collected and aliquoted. It was stored at −20 °C until analysis. The Pierce™ BCA Protein Assay Kit (Thermo Scientific™; #23227) was used to measure protein concentration. Protein separation was performed using a concentration of 10 μg/μL of 4-15% Mini-PROTEAN® TGX™ Precast Protein Gels, 10-well, 50 µL (Bio-Rad; #4561084). Protein transfer was performed by wet blotting onto a PVDF membrane (Life Technologies, Carlsbad, CA, USA). The blots were blocked with 5% bovine serum albumin (Sigma-Aldrich; *#*05470) or 5% non-fat dry milk (Cell Signaling Technology; #9999) in Tris-buffered saline/Tween-20 (TBST), depending on the requirements of the antibody. The membranes were then incubated with primary antibodies in blocking solution overnight at 4°C, washed with TBST and then incubated with secondary antibodies in TBST. Antibodies against the specific proteins used are listed in Table 1. To visualize the bands, a peroxidase goat antibody was used as a secondary antibody against the host of each primary antibody (J. ImmunoResearch). All secondary antibodies were used at a concentration of 1:10,000. Secondary antibodies were conjugated to horseradish peroxidase (HRP) and protein bands were visualised by chemiluminescence using Clarify™ Western ECL Substrate (Bio-Rad) and imaged using Azure Biosystems c600. Relative signal intensity was quantified using Fiji software and normalised to the total protein content of housekeeping protein and/or uninjured-uninjected control^57^. The experiments were replicated by two independent experimenters.

#### mRNA isolation and qRT-PCR

RNA was isolated using the RNeasy Micro Kit (QIAGEN; #74004) according to the manufacturer’s protocol. A NanoPhotometer© P330 (Implen, Munich, Germany) was used to quantify the amount of isolated RNA. RNA was then reverse transcribed into complementary DNA (cDNA) using Transcriptor Universal cDNA Master (Roche; #05893151001) in a T100TM thermal cycler (Bio-Rad, Hercules, CA, USA) according to the manufacturer’s protocol. For qPCR, TaqMan® gene expression assays (Life Technologies by Thermo Fisher Scientific, Waltham, MA, USA) were used for CREB1 (Rn00578828_g1), Itga9 (Rn01746751_m1) and GAPDH (Rn01775763_g1), all purchased from Applied Biosystems and used according to the manufacturer’s recommendations. FastStart Universal Probe Master (Rox) (Roche; #FSUPMMRO) was used as the master mix. The amplification was performed on a qRT-PCR cycler (QuantStudio® 6 Flex PCR System, Applied Biosystems® from Thermo Fischer Scientific). For all amplifications, the same cycling conditions were used: 2 min at 50°C, 10 min at 95°C, followed by 40 cycles of 15 s at 95°C and 1 min at 60°C. The threshold cycle value (Ct) was used to calculate gene expression. The expression was normalised to the uninjected and uninjured control. Each of the qRT-PCR experiments was carried out in duplicate. The Ct values of each measurement condition were normalised to GAPDH. Values were then expressed as log_2_ fold changes according to Livak and Schmittgen^58^. The experiments were replicated by two independent experimenters.

#### Statistical analysis

Statistical analysis was performed using GraphPad Prism 10, and statistical differences between groups were determined using t-test, one-way or two-way ANOVA, with appropriate post hoc tests when necessary. Data were not tested for normality. No outlier test was performed. A p-value of 0.05 was considered significant for all statistical analyses. The statistical tests and appropriate post hoc tests used in each analysis are also indicated in the results sections of the respective figures.

## Acknowledgment

We would like to thank Prof. James Fawcett from the University of Cambridge for all his valuable advice. Microscopy was done at the Microscopy Service Centre of the Institute of Experimental Medicine CAS supported by the MEYS CR (LM2023050 Czech-Bioimaging). We also acknowledge the Light Microscopy Core Facility, IMG, Prague, Czech Republic, supported by MEYS (LM2018129, CZ.02.1.01/0.0/0.0/18_046/0016045) and RVO: 68378050-KAV-NPUI, for their support with the Light sheet fluorescence imaging and image analysis presented herein. This work was supported by Czech Science Foundation (24-11193S); Ministry of Education, Youth and Sports (Excellence in Regenerative Medicine; CZ.02.01.01/00/22_008/0004562); and by Charles University Grant Agency, project No. 320421 and 102122.

## Author Contributions

KS, AC, and LMU designed the research and interpreted the data; KS, AC, BS, RH, LB, JC, VS and DM performed research; KS, AC, LB, JC analysed data; JvdH and FdW prepared the AAV vectors; KS, BS and PJ obtained the funding. All authors were involved in manuscript writing and revision.

## Data availability statement

Access to data is available in Zenodo (10.5281/zenodo.14196257) upon request.

## Ethics approval statement

All procedures were approved by the Ethics Committee of the Institute of Experimental Medicine of the ASCR and were performed in accordance with Act No. 77/2004 of the Czech Republic (ethics approval number: AVCR 4386/2022 SOV II).

